# Robust Estimation of Noise for Electromagnetic Brain Imaging with the Champagne Algorithm

**DOI:** 10.1101/2020.08.07.241943

**Authors:** Chang Cai, Mithun Diwakar, Ali Hashemi, Stefan Haufe, Kensuke Sekihara, Srikantan S Nagarajan

**Affiliations:** Department of Radiology and Biomedical Imaging, University of California, San Francisco, CA 94143-0628; Berlin Center for Advanced Neuroimaging, Charité Universitätsmedizin Berlin, Berlin, Germany; Machine Learning Group, Electrical Engineering and Computer Science Faculty, Technische Universität Berlin, Germany; Institut für Mathematik, Technische Universität Berlin, Germany; Bernstein Center for Computational Neuroscience Berlin, Berlin, Germany; Department of Advanced Technology in Medicine, Tokyo Medical and Dental University, 1-5-45 Yushima, Bunkyo-ku, Tokyo 113-8519, Japan; Signal Analysis Inc., Hachioji, Tokyo

**Keywords:** Electromagnetic Brain Mapping, Robust Noise Estimation, Bayesian Inference, Inverse problem, Magnetoencephalography

## Abstract

Robust estimation of the number, location, and activity of multiple correlated brain sources has long been a challenging task in electromagnetic brain imaging from M/EEG data, one that is significantly impacted by interference from spontaneous brain activity, sensor noise, and other sources of artifacts. Recently, we introduced the Champagne algorithm, a novel Bayesian inference algorithm that has shown tremendous success in M/EEG source reconstruction. Inherent to Champagne and most other related Bayesian reconstruction algorithms is the assumption that the noise covariance in sensor data can be estimated from “baseline” or “control” measurements. However, in many scenarios, such baseline data is not available, or is unreliable, and it is unclear how best to estimate the noise covariance. In this technical note, we propose several robust methods to estimate the contributions to sensors from noise arising from outside the brain without the need for additional baseline measurements. The incorporation of these methods for noise covariance estimation improves the robust reconstruction of complex brain source activity under high levels of noise and interference, while maintaining the performance features of Champagne. Specifically, we show that the resulting algorithm, Champagne with noise learning, is quite robust to initialization and is computationally efficient. In simulations, performance of the proposed noise learning algorithm is consistently superior to Champagne without noise learning. We also demonstrate that, even without the use of any baseline data, Champagne with noise learning is able to reconstruct complex brain activity with just a few trials or even a single trial, demonstrating significant improvements in source reconstruction for electromagnetic brain imaging.

## 1. Introduction

Electromagnetic brain imaging is the process of measuring and spatio-temporal reconstruction of brain activity from non-invasive sensor recordings of magnetic fields and electric potentials. In order to transform these sensor recordings into brain activity, both the forward and inverse models must be solved. The forward model combines source, volume conductor, and measurement models to estimate a mixing matrix called the leadfield that describes a linear relationship between sources and the measurements. Solving the inverse problem is the process of devising inverse algorithms to estimate the parameters of brain activity from sensor data and the leadfield matrix.

Most inverse algorithms can be viewed in a Bayesian framework [1, 2]. This perspective is useful because at a high level, the prior distribution, implicitly or explicitly imposed, can be used to differentiate and compare the various source localization methods. Recently, we developed Champagne, a novel tomographic source reconstruction algorithm derived in an empirical Bayesian fashion with incorporation of deep theoretical ideas about sparse-source recovery from noisy, constrained measurements. Champagne improves upon existing methods of source reconstruction in terms of reconstruction accuracy, robustness, and computational efficiency [3]. Experiments with preliminary simulated and real data, presented in [4], show that compared to other commonly-used source localization algorithms, Champagne is more robust to interference from correlated sources and noisy data.

Inherent to Champagne is the availability of “baseline” data to estimate the statistics of the “brain noise”. However, in many MEG and EEG experimental scenarios, “baseline” data is not available or is unreliable. For example, in certain paradigms, there are artifacts that are only present in the “active” time-period and absent in the baseline or control period, such as during experiments with speaking or other muscle movements. Also, in experiments involved event-related desynchronization, the signal amplitude decreases relative to the baseline levels. In such situations, it is unclear how noise estimation should be accomplished for algorithms such as Champagne. This predicament also applies to other Bayesian source reconstruction methods such as Saketini [5], NSEFALoc [6] and LowSNR-BSI [7].

In this technical note, we propose robust methods to estimate the contributions to sensors from noise without the need for additional “baseline” or “control” measurements. Importantly, the proposed methods preserve the robust reconstruction performance features of the sparse source reconstruction algorithm Champagne. Our novel robust noise estimation algorithms partition contributions to the sensor data from brain activity sources and noise related activity, with corresponding Gaussian variance parameters for both brain sources and noise that are estimated from data. Variance parameters are estimated using empirical Bayesian inference, i.e. maximizing the marginal likelihood of the data.

The resulting inference algorithms are quite robust to the prior initialization variance, to different noise modalities, to the reconstruction of highly correlated multiple sources, and to the effect of high levels of interference and noise. In simulations, performance of the proposed noise learning algorithms are consistently superior to original Champagne without noise learning. Without baseline data, the novel noise learning algorithms are robust to correlated brain activity present in real data sets and are able to reconstruct complex brain activity with few trials or even a single trial, demonstrating significant improvements in electromagnetic brain imaging.

Section 2 describes the original Champagne algorithm and Champagne algorithm with noise learning in detail. Section 3 describes the tested simulation configurations, performance evaluation metrics, and analyzed real datasets. Section 4 describes performance results in simulated and real data, followed by brief discussion in Section 5.

## 2. Methods

This section first briefly reviews the probabilistic generative model for electromagnetic brain imaging used by the Champagne algorithm, the original Champagne algorithm which provides the necessary update rules for brain sources time course and voxel variance estimates with the statistics of the background activity estimated from “baseline” or “control” measurements. Following this, we propose robust methods to estimate the contributions to sensors from noise arising from outside the brain without the need for additional baseline measurements. In summary, the Champagne algorithm with noise learning initializes the voxel variances and noise covariance and updates them until the marginal likelihood converges, and after convergence outputs the brain source activity time-courses.

### 2.1. Probabilistic generative model for electromagnetic brain imaging

We assume that MEG or EEG data have been collected for induced or spontaneous brain activity paradigms, with separate time-windows for induced or spontaneous source activity and for background interference including biological, environmental sources, and sensor noise. We define the following parameters: ***y***(*t*) ∈ ℝ^*M ×*1^, is the output data of sensors at time *t, M* is the number of channels measured. ***L*** = [***L***_1_, …, ***L***_*N*_] is the leadfield matrix from the forward model. *N* is the number of voxels under consideration and 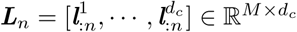 is the leadfield matrix for *n*-th voxel. The *k*-th column of ***L***_*n*_ represents the signal vector that would be observed at the scalp given a unit current source or dipole at the *n*-th voxel with a fixed orientation in the *k*-th direction. The voxel dimension *d*_*c*_ is usually set to 3. Multiple methods based on the physical properties of the brain and Maxwell’s equations are available for the computation of each ***L***_*n*_ [8]. For convenience, we also define ***L*** = [***l***_1:_, …, ***l***_*M*:_]^*T*^, where ***l***_*m*:_ denotes the *m*-th row vector of ***L***. 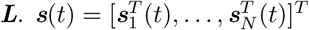 is the unknown brain activity. 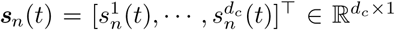 is the *n*-th voxel intensity at time *t*, which we assume it with *d*_*c*_ orientations. The generative probabilistic model for the sensor data at time point *t* can be written as:

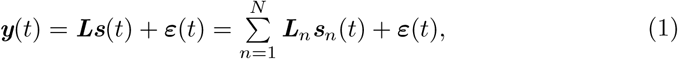

with prior distributions ***s***(*t*) ∼ 𝒩 (***s***(*t*)|***0, α***) and ***ε***(*t*) ∼ 𝒩 (0, **Λ**). Where **Λ** = diag(*λ*_1_, *λ*_2_, …, *λ*_*M*_) and ***α*** is defined as *d*_*c*_*N* × *d*_*c*_*N* block diagonal matrix expressed as

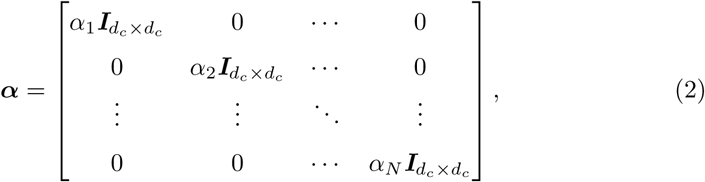

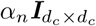 is a prior variance *d*_*c*_ × *d*_*c*_ matrix of ***s***_*n*_ and 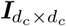 is a *d*_*c*_ × *d*_*c*_ identity For simplicity, we define matrix ***Y*** = [***y***(1), …, ***y***(*T*)] and ***S*** = [***s***(1), …, ***s***(*T*)] as the entire sensor and source time series, where *T* is the number of time points. The prior distribution *p*(***S***|***α***) is then defined as

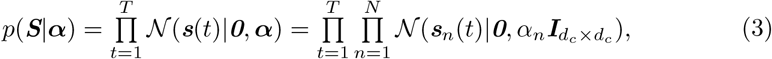

Our goal is to jointly estimate the posterior mean 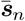, the parameters ***α*** and **Λ**.

### 2.2. The original Champagne algorithm

We provide a brief overview of the original Champagne algorithm, detailed derivations can be found in [3, 9]. Important to note that, in the original Champagne algorithm, the noise covariance **Λ** is learnt from available baseline or control measurements [3]. In contrast, here we describe update rules for the noise covariance estimation without baseline measurements.

Brain source activity is estimated from the posterior distribution of the voxel activity *p*(***S***|***Y***). The voxel variance hyperparameters are estimated by maximizing a bound on the marginal likelihood *p*(***Y***|***α***). Although there are multiple ways to derive update rules for ***α*** [2], in Champagne we utilize a convex bounding [10] on the logarithm of marginal likelihood (model evidence), which results in fast and convergent update rules [11]. For a detailed derivation of the original Champagne, we refer to our previous paper [3]. **Table 1** lists the sources level updates and the cost function used in Champagne.

**Table 1.**
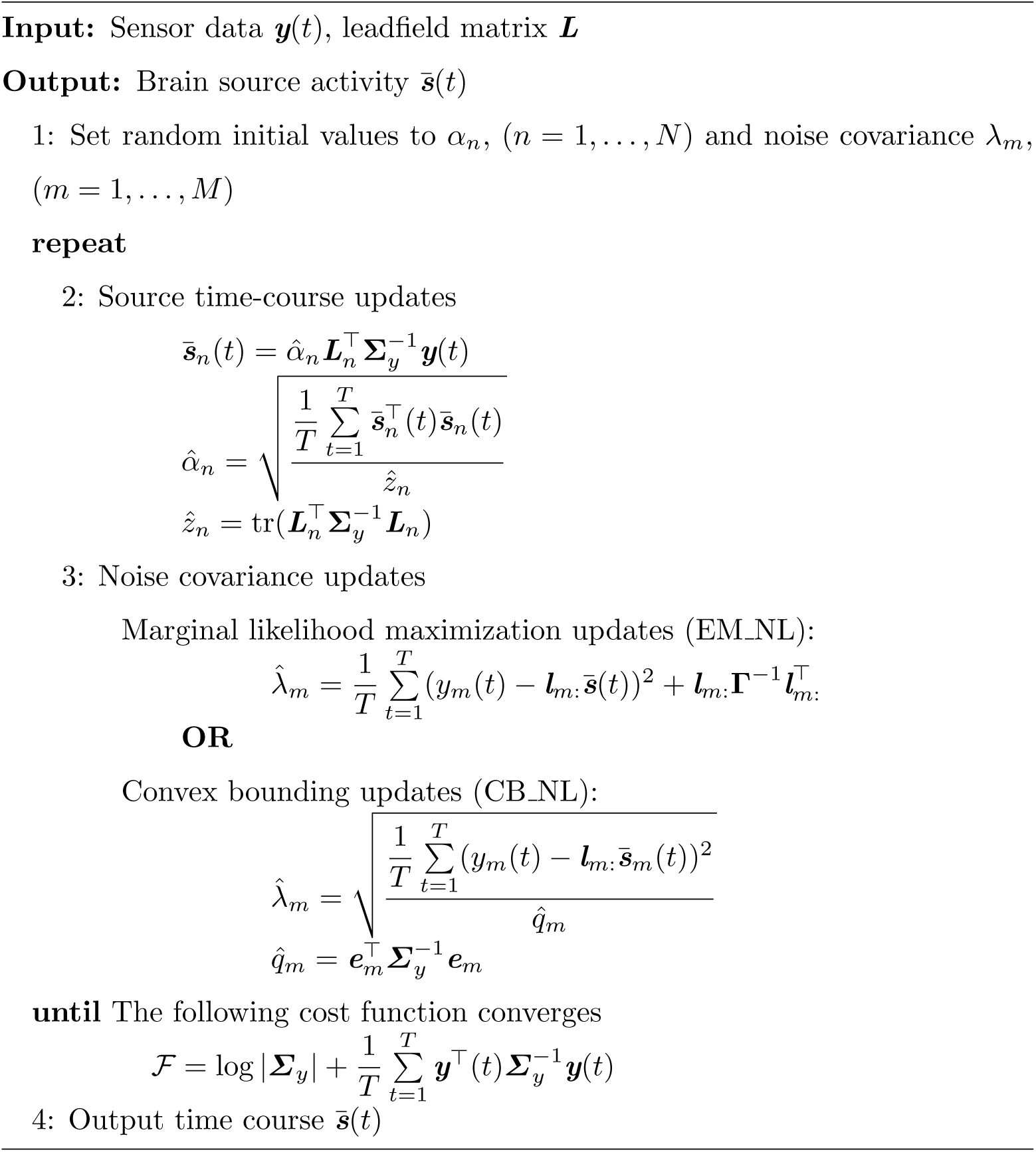
The Champagne Algorithm with Noise Learning.

### 2.3. Noise covariance estimation (**Λ**)

The diagonal noise covariance can be defined with two different structures: homoscedastic and heteroscedastic [12, 13]. Homoscedastic noise assumes the noise variance is the same for all sensors. In contrast, heteroscedastic noise assumes the noise variance differs between sensors. In this paper, we assume the noise covariance is heteroscedastic, as homoscedastic noise can be easily handled by computing a scalar average of the diagonal elements of the heteroscedastic covariance matrix and multiplying by an *M* × *M* identity matrix. We introduce three ways to derive the update rules for the noise covariance **Λ**: marginal likelihood maximization, expectation-maximization, and convex-bounding of the marginal likelihood.

#### 2.3.1. Learning noise covariance using marginal likelihood maximization

The noise covariance **Λ** can be directly estimated by setting the derivative of the cost function with respect to **Λ** to zero with fixed ***α***, more details is shown in Appendix A, the update rule for *λ*_*m*_ is expressed as:

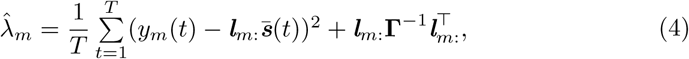

where **Γ** = ***α***^−1^ + ***L*^*T*^Λ^−1^*L***. An alternative derivation of the noise covariance using the Expectation Maximization (EM) algorithm is shown in Appendix B which results in identical update rules.

#### 2.3.2. Learning noise covariance by maximizing convex bound of marginal likelihood

Another way to estimate the noise covariance is using an auxiliary cost function which is based on maximizing a convex bounding function of the marginal likelihood. This method has the following advantages: first, the convex bounding approach is guaranteed to reduce the cost function at each iteration; second, it shows higher computation efficiency and faster convergence compared to EM-based estimation; third, noise learning and voxel variance learning can be easily unified to produce one simple generative model. Noise covariance update rules can be derived through convex bounding of the cost function, more details is shown in Appendix C, the update rules are:

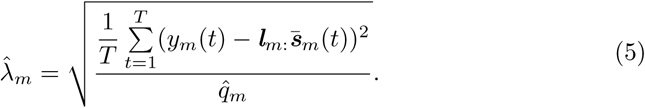

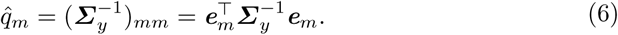

where ***q*** = [*q*_1_, *q*_2_, …, *q*_*M*_]^*T*^ is an auxiliary variable, ***e***_*m*_ is a *M* × 1 vector and the element in *m*-th row is 1, the others are 0. With convex-bounding (CB) based update rules for noise, it is possible to efficiently estimate the voxel and noise covariance simultaneously by lead-field augmentation, as described in Appendix D.

### 2.4. Summary

In summary, as is shown in **Table 1**, the Champagne algorithm with noise learning does not need the input of noise covariance **Λ**, it first initializes the voxel variances and noise covariance with random values, then updates the voxel variances (the same as the original Champagne) and the noise covariance using EM or CB. Each iteration of this Champagne algorithm with noise learning theoretically guarantees to reduce (or leave unchanged) the cost function of the data. Finally, the Champagne algorithm with noise learning outputs the brain activity time courses.

## 3. Performance Evaluation of Simulated and Real Data

### 3.1. Benchmarks for Comparison

We implemented two variants of Champagne with noise learning - 1) using marginal likelihood maximization or expectation maximization (EM) and 2) using convex bounding (CB) which are referred to as EM_NL and CB_NL respectively. For simplicity, this paper does not show the performance results of noise-learning with homoscedastic noise as its performance is comparable to the use of heteroscedastic noise. We compare these two variants of Champagne with noise learning with three different benchmarks. A first benchmark we use is the original Champagne with different fixed levels of noise estimated by noise subspace decomposition of the sample data covariance. We refer to this algorithm as Champagne with Noise_Sub. A second benchmark is the original Champagne where we use available baseline data (in simulations) to learn a low-rank non-diagonal noise covariance using the Variational Bayes Factor Analysis algorithm (VBFA) [14]. This benchmark would represent an upper bound on the performance of Champagne with noise learning when baseline data is available. For real data, we also include sLORETA as a third benchmark algorithm for comparison.

### 3.2. Simulation configurations

We generate data by simulating dipole sources with fixed orientation. Damped sinusoidal time courses with frequencies sampled randomly between 1-75Hz are created as voxel source time activity and then projected to the sensors using the leadfield matrix generated by the forward model. We assume 271 MEG sensors and a single-shell spherical model [8] as implemented in SPM12 (http://www.fil.ion.ucl.ac.uk/spm) at the default spatial resolution of 8196 voxels at approximately 5 mm spacing. The time period is set as 480 samples with source activities of interest and noise activity. To evaluate the robustness of the proposed noise learning methods, we randomly choose noise activity with real brain noise consisting of actual resting-state sensor recordings collected from ten human subjects presumed to have only spontaneous brain activity and sensor noise. Signal-to-noise ratio (SNR) and correlations between voxel time courses are varied to examine algorithm performance. SNR and time course correlation are defined in a standard fashion [4, 15].

We examined performance for reconstruction of 5 random seeded point sources with SNR of 3 dB (real brain noise) and inter-source correlation coefficient of 0.99. The ratio of noise covariance to sample data covariance for all algorithms was increased from 0.05% to 10%.

We increased the number of seeded point sources to evaluate the algorithm performance as a function of the number of sources. Inter-source correlation coefficient was fixed at 0.99 and SNR was fixed at 3 dB. The number of seeded point sources was increased from 3 to 15 with a step of 2.

We evaluated algorithm performance as a function of SNR. Reconstruction performance was evaluated for 5 randomly seeded point sources with an intersource correlation coefficient of 0.99. Simulations were performed at SNRs from −10 dB to 20 dB in steps of 5 dB.

For Champagne with Noise_Sub, the noise covariance **Λ** is assumed to be fixed as a scalar value multiplied by an identity matrix, and computed as a percentage of the norm of the sample data covariance.

### 3.3. Real datasets

Real MEG data was acquired in the Biomagnetic Imaging Laboratory at University of California, San Francisco (UCSF) with a CTF Omega 2000 whole-head MEG system from VSM MedTech (Coquitlam, BC, Canada) with 1200 Hz sampling rate. The leadfield for each subject was calculated in NUTMEG [16] using a single-sphere head model (two spherical orientation leadfield) and an 8 mm voxel grid. Each column was normalized to have a norm of unity. The data were digitally filtered from 1 to 70 Hz to remove artifacts and DC offset.

Three real MEG data sets were used to evaluate the application of the algorithms: 1. Somatosensory Evoked Fields (SEF); 2. Auditory Evoked Fields (AEF); 3. Resting state data. The first two data sets have been reported in our prior publications using the Champagne algorithm, and details about these datasets can be found in [3, 4]. In order to evaluate the robustness of our novel algorithms to scarcity of data, we collected SEF and AEF data from five subjects (around 250 trials per subject for SEF, 120 trials per subject for AEF). We tested reconstruction with the number of trials limited to 1, 2, 12 and 63 for both SEF and AEF. Each reconstruction was performed 30 times with the specific trials themselves chosen as a random subset of all available trials. For resting state data analysis, six subjects were instructed simply to keep their eyes closed and their thoughts clear. We collected 4 trials per subject, each trial of 1 minute length with a sampling rate of 1.2 kHz. We randomly chose 10 seconds or equivalently 12,000 time samples for brain source reconstruction from each subject.

### 3.4. Quantifying algorithm performance

To evaluate the performance of localization results, we use free-response receiver operator characteristics (FROC) which shows the probability for detection of a true source in an image versus the expected value of the number of false positive detections per image [4, 17, 15, 18]. Based on the FROC, we compute an *A*′ metric [19, 20] which is an estimate of the area under the FROC curve for each simulation. If the area under the FROC curve is large, then the hit rate is higher compared to the false positive rate. The *A*′ metric is computed as follows:

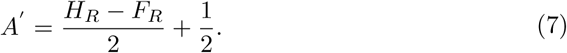

Hit rate (*H*_*R*_) is calculated by dividing the number of hit point sources by the true number of point sources. The false rate (*F*_*R*_) is defined by dividing the number of potential false positive voxels by the total number of false voxels for each simulation. The details of the *A*′ metric calculation can be referred to in our previous paper [20]. We then calculate the correlation coefficient between the seed and estimated source time courses for each hit, which is used to determine the accuracy of the time course reconstructions and denoted as 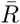. Finally, we combine these two metrics that capture both the accuracy of the location and time courses of the algorithms into a single metric called the Aggregate Performance (AP) [4, 15, 20]:

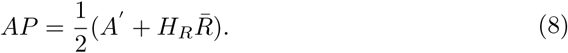

The AP ranges from 0 to 1, with higher numbers reflecting better performance [4, 17, 15, 18]. To calculate the mean and variance of AP, the results were averaged with 50 simulations at each configuration.

### 3.5. Algorithm initialization

The initialization for the algorithms are as follows. For Champagne with Noise_Sub, the noise covariance **Λ** is assumed to be fixed as a scalar value multiplied by an identity matrix, and computed as a percentage of the norm of the sample data covariance. A range percentages from 0.05% to 10% are utilized. For both variants of Champagne with noise learning, the initialization for noise covariance is set to the equivalent Noise_Sub covariance matrix. Initialization of hyperparameters for brain source activity ***α*** is performed by computing the Minimum-Norm Estimation (MNE) [1] to estimate voxel variances. For the benchmark algorithm sLORETA used in real datasets, we use the default setting in NUTMEG software where the regularization parameter is equal to the maximum eigenvalue of the sensor data covariance [16]. In real data sets, since we do not know the details of the noise, ground truth is defined as the brain activity estimated from all available trials for each subject.

## 4. Results

### 4.1. Simulation results

In Figure 1, five random point sources are seeded with inter-source correlation coefficient of 0.99. The SNR is set as 3 dB with real brain noise. The ratio of noise covariance to sample data covariance for all algorithms are increased from 0.05% to 10%. As shown, EM_NL and CB_NL show similar aggregate performance results, which consistently outperform the Champagne algorithms without noise learning and are close to original Chamapagne with true non-diagonal noise. In contrast, increasing the ratio decreases the performance of Champagne with Noise_Sub.

**Figure 1:**
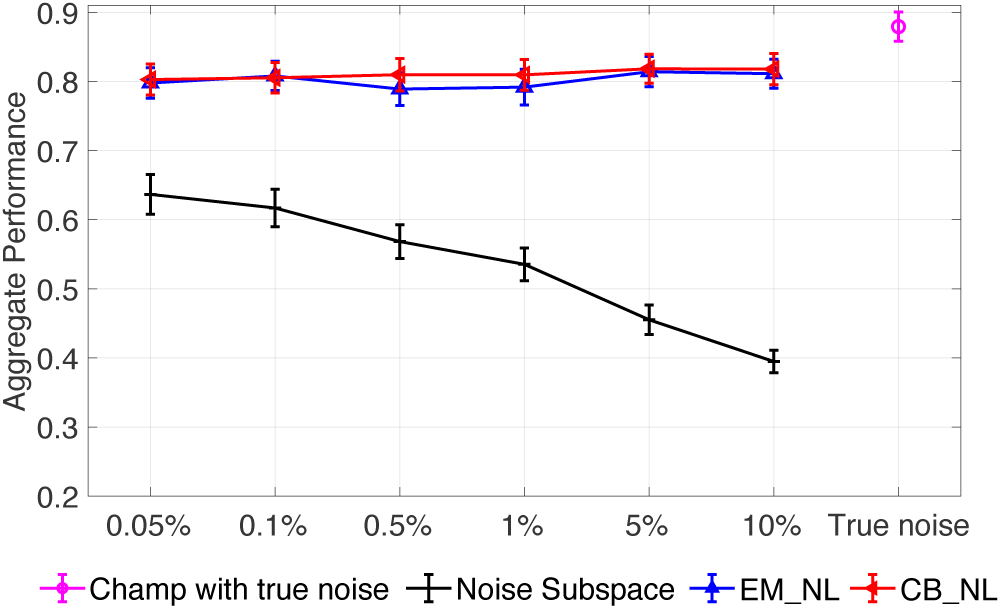
Aggregate performance in simulations for different noise levels for Champagne with true non-diagonal noise, Noise_Sub and for the noise learning algorithms proposed here. Five random point sources are seeded with inter-source correlation coefficient of 0.99. The SNR is set to 3 dB with real brain noise. Results are averaged with 50 simulations at each data point and the error bars show the standard error.

Figure 2 plots aggregate performance results of varying the number of seeded point sources and SNR. Results of all algorithms in response to increasing number of seeded point sources are presented in Figure 2(A). Here, the inter-source correlation coefficient is fixed at 0.99 and SNR is fixed at 3 dB. All algorithms have the same trend, showing decreasing performance as number of sources increases. In general, the performance of EM_NL and CB_NL are similar and close to original Champagne with true non-diagonal noise covariance setting. In addition, EM_NL and CB_NL consistently show better performance results than original Champagne with Noise_Sub. When increasing the ratio of noise subspace covariance to data covariance, the performance of Champagne with Noise_Sub decreases. Champagne with Noise_Sub (0.1%) produces the best performance among all noise subspace covariance settings and Champagne with Noise_Sub (10%) produces the worst performance.

**Figure 2:**
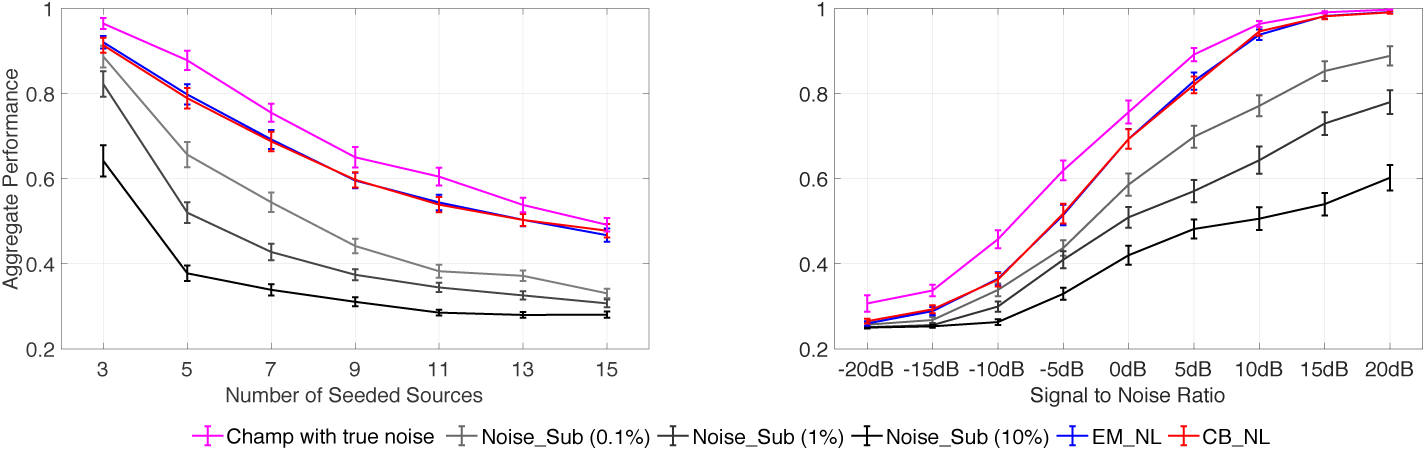
Simulation results of aggregate performance versus (A) number of seeded point sources; (B) signal-to-noise ratio. The results are averaged with 50 simulations at each data point and the error bars show the standard error.

Performance results versus SNR for all algorithms are plotted in Figure 2(B). Reconstruction performance is evaluated for five randomly seeded point sources with an inter-source correlation coefficient of 0.99. Again, all algorithms have the same trend, with increasing performance as the SNR increases. EM_NL and CB_NL perform similar to original Champagne with true non-diagonal noise covariance setting and consistently produce more accurate results than Champagne with Noise_Sub. Champagne with Noise_Sub (0.1%) produces the best performance among all non-noise learning algorithms and Champagne with Noise_Sub (10%) shows the worst performance.

In summary, from the results of our computer simulations, we can conclude that EM_NL and CB_NL consistently show similar and closer performance to original Champagne with true non-diagonal noise, which outperform Champagne with Noise_Sub (0.1% to 10%). Incorrect noise covariance estimation for traditional Champagne generates poor performance, as expected. Since EM_NL and CB_NL show similar performance, for simplicity, in the next section for real data sets, we only present the performance of CB_NL.

### 4.2. Results of real datasets

#### 4.2.1. Somatosensory evoked field paradigm

Figure 3 shows localization results of the somatosensory evoked field response due to somatosensory stimuli presented to one representative subject’s right index finger. A peak should be typically seen ∼50 ms after stimulation in the contralateral (in this case, the left) somatosensory cortical area for the hand, i.e., dorsal region of the postcentral gyrus (hand knob). To evaluate algorithmic robustness, we randomly choose several subsets of trials from the full 252 trials for reconstruction. Since for SEF data sets, there is only one brain source, we present another widely used benchmark for comparison-sLORETA. As is shown, Champagne with CB_NL is able to localize activation to the correct area of the somatosensory cortex with focal reconstructions under even a few trials or a single trial. Champagne with Noise_Sub (10%) is able to produce reasonable results when the number of trials is equal or larger than 12, otherwise, localization with Champagne with Noise_Sub is biased towards the edge of the head and contains several false areas of brain activity. sLORETA is able to localize diffuse brain activity at the somatosensory cortex with 12 and 63 trials; otherwise, the activity is falsely localized.

**Figure 3:**
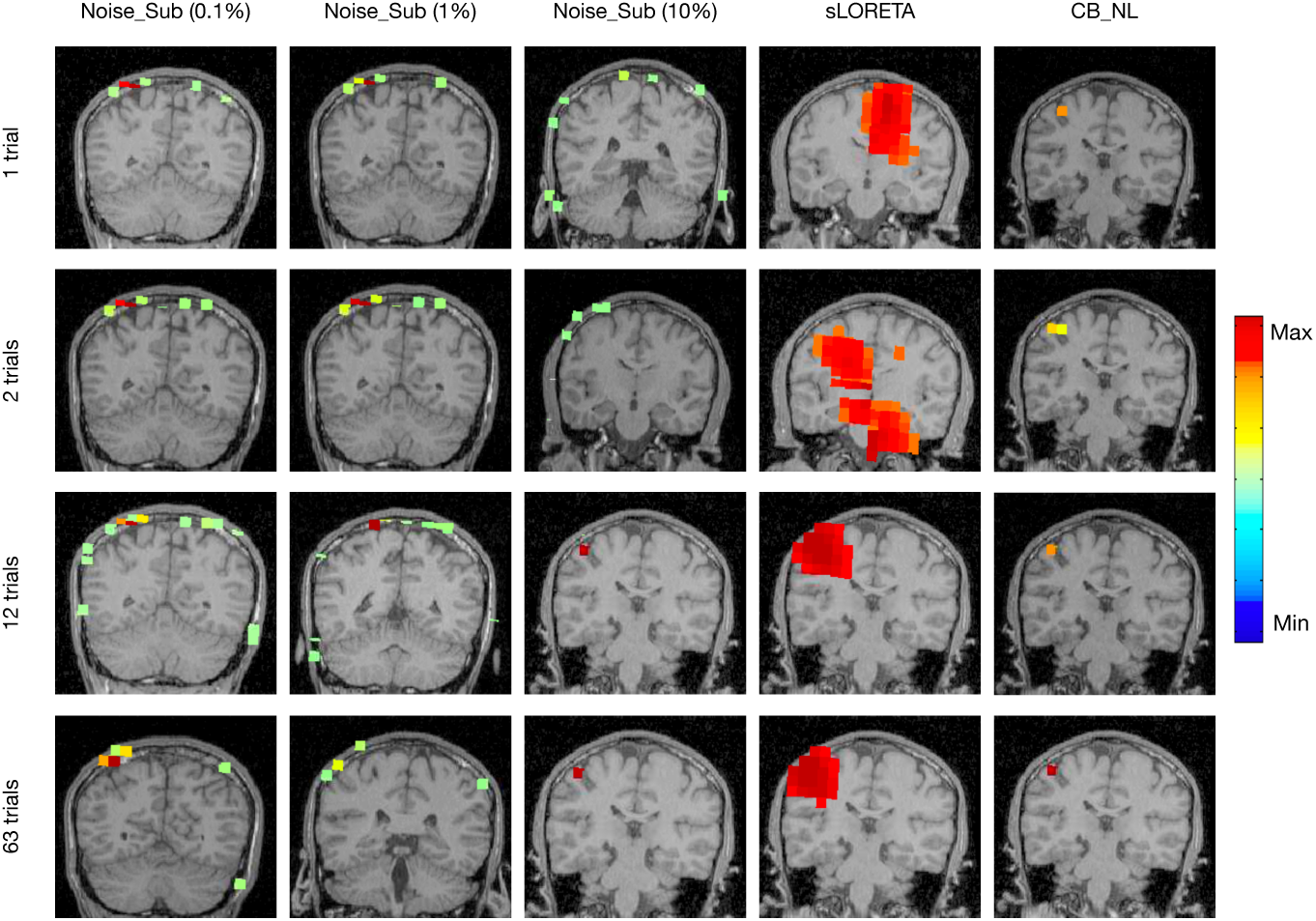
Sensory Evoked Field localization results versus number of trials using Champagne with Noise_Sub (0.1% to 10%), sLORETA, and Champagne with CB_NL.

Figure 4 shows five individuals and averaged aggregate performance results of Champagne with Noise_Sub (0.1% to 10%), sLORETA, and Champagne with CB_NL for sensory evoked field localization versus number of trials. Error bars depict standard errors. Trials are randomly chosen from around 250 trials from each subject and the number of trials is increased from 1 to 63. Each condition is tested 30 times for each subject. Ground truth is defined as the brain activity estimated from around 250 trials per subject. The same strategy is used as for simulations to obtain the aggregate performance for each test. In general, increasing the number of trials increases the performance of all algorithms. Champagne with CB_NL consistently produces better results than Champagne with Noise_Sub and sLORETA.

**Figure 4:**
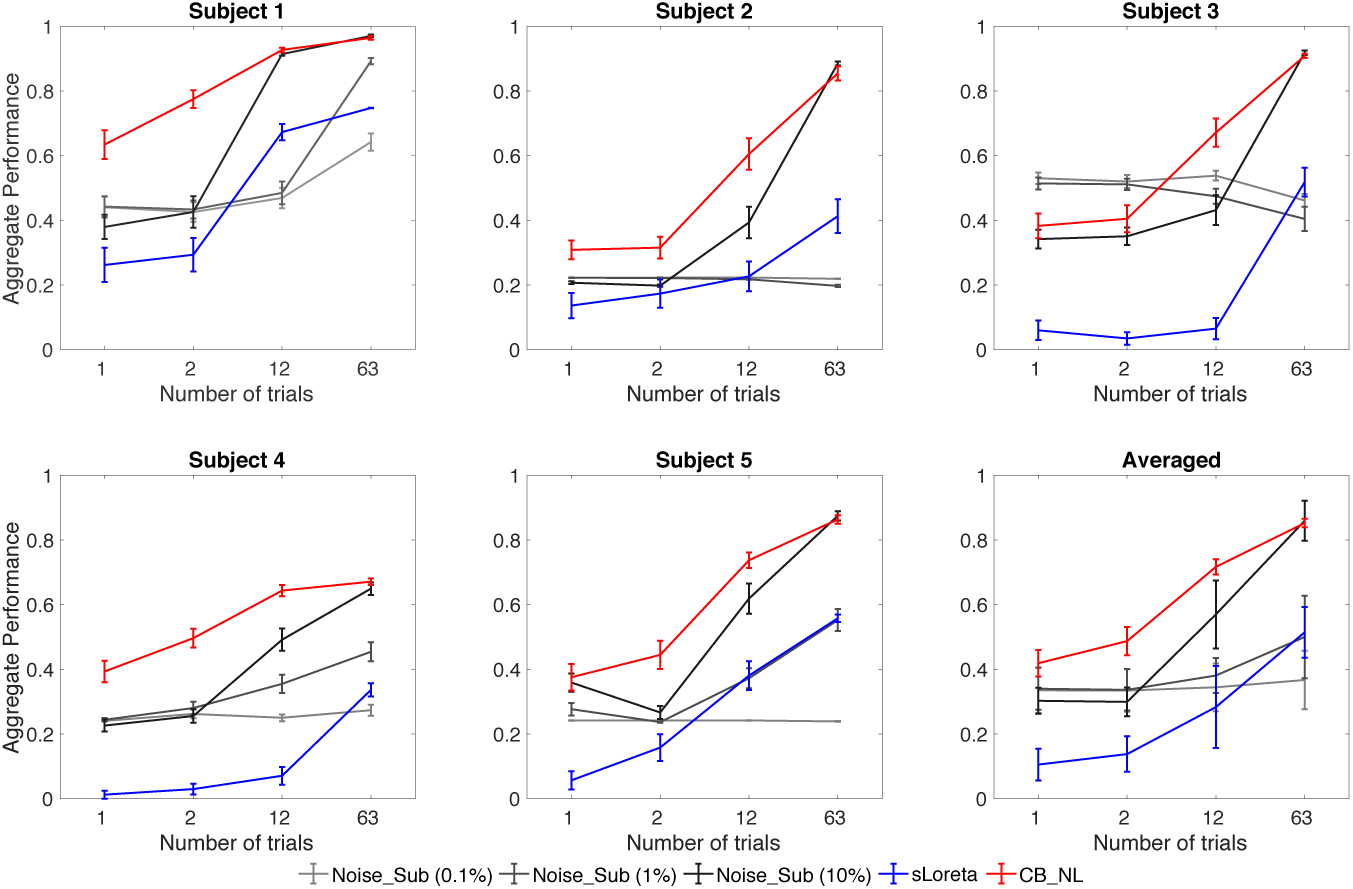
Aggregate performance results versus number of trials for SEF for five subjects using Champagne with Noise_Sub (0.1% to 10%), sLORETA, and Champagne with CB_NL.

Figure 5 shows Auditory evoked field (AEF) localization results versus number of trials from a single representative subject using Champagne with Noise_Sub, sLORETA, and Champagne with CB_NL. The power at each voxel around the M100 peak is plotted for each algorithm. Again, Champagne with CB_NL is able to localize the expected bilateral brain activation with focal reconstructions under even a few trials or even a single trial. The limited number of trials does not influence the reconstruction results. Specifically, the activities localize to Heschl’s gyrus in the temporal lobe, which is the characteristic location of the primary auditory cortex.

**Figure 5:**
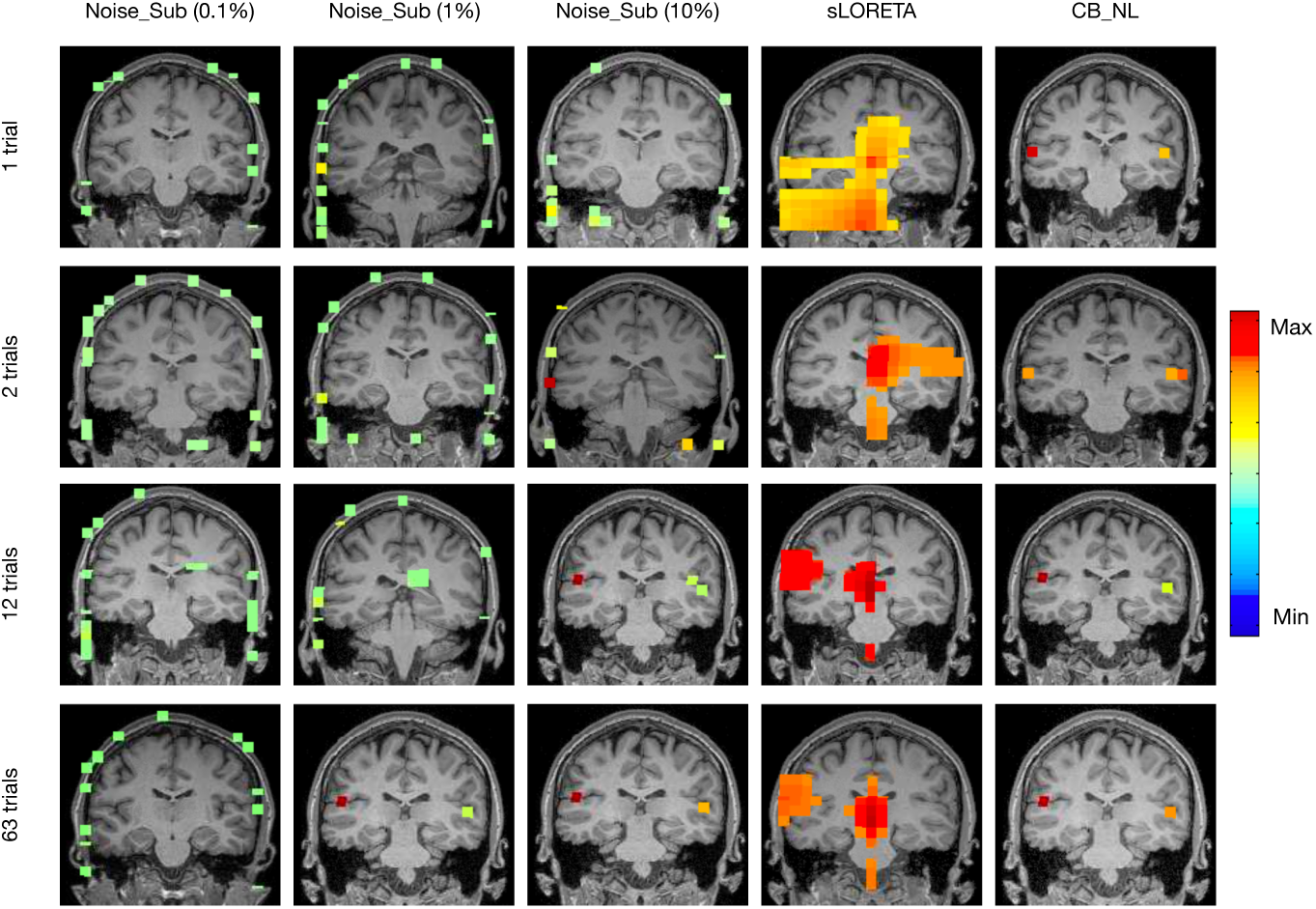
Auditory evoked field (AEF) localization results versus number of trials from one representative subject using Champagne with Noise_Sub (0.01% to 10%), sLORETA, and Champagne with CB_NL.

Champagne with Noise_Sub (10%) is able to localize the bilateral auditory activity when the number of trials is larger than 12 and Champagne with Noise_Sub (1%) is able to localize the bilateral auditory activity with the number of trials is 63; otherwise, localization by Champagne with Noise_Sub is biased towards the edge of the head and produces several areas of pseudo brain activity. Inaccurate noise covariance estimation further degrades the performance of Champagne with Noise_Sub. sLORETA is unable to localize bilateral auditory activity in all conditions.

Figure 6 shows five individuals and averaged aggregate performance results of Champagne with Noise_Sub (0.1% to 10%), sLORETA, and Champagne with CB_NL for Auditory Evoked Field localization versus number of trials; the error bars show standard error. Trials are randomly chosen from around 120 trials from each subject and the number of trials is increased from 1 to 63. Each condition is tested 30 times for each subject. Again, the ground truth is defined as the brain activity estimated from around 120 trials per subject. In general, increasing the number of trials increases the performance of all algorithms. Champagne with CB_NL consistently produces more accurate results than Champagne with Noise_Sub and sLORETA.

**Figure 6:**
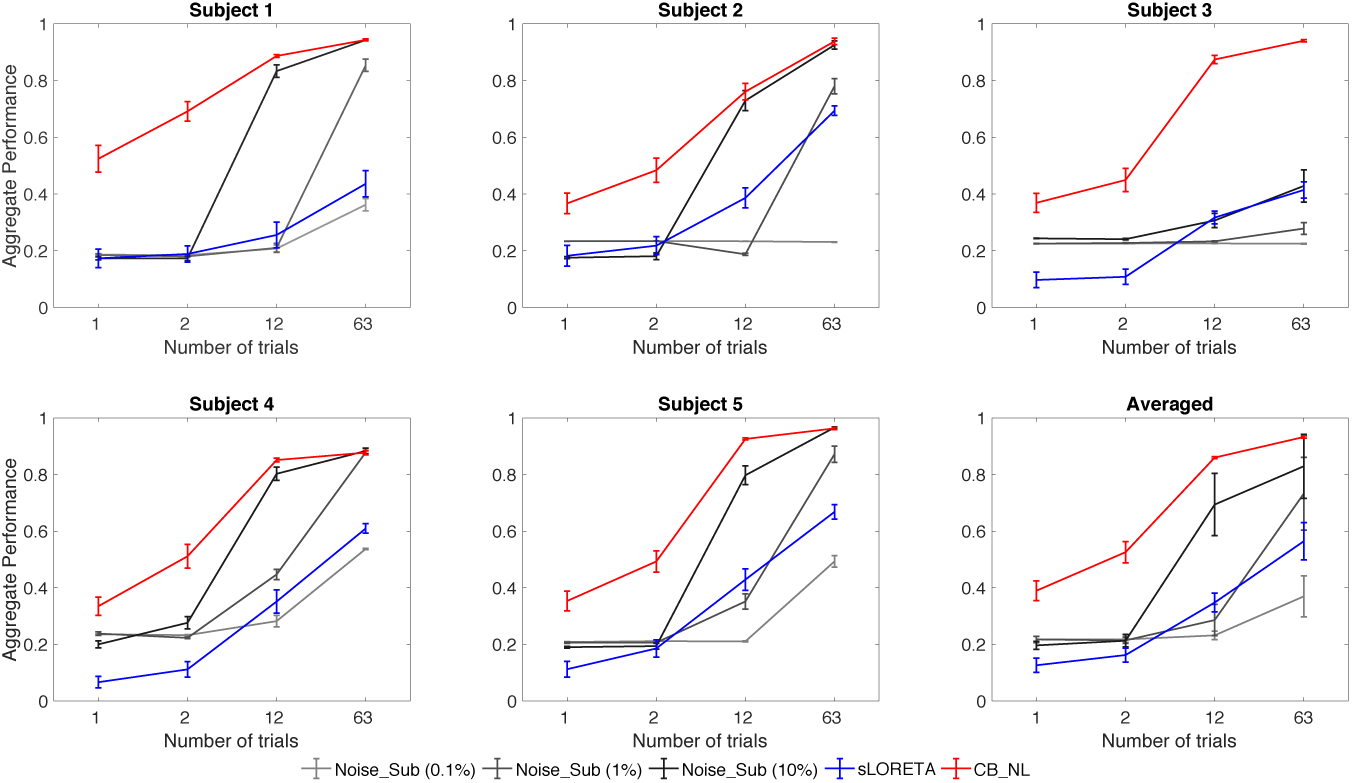
Aggregate performance results versus number of trials for AEF from five subjects using Champagne with Noise_Sub (0.01% to 10%), sLORETA, and Champagne with CB_NL.

#### 4.2.3. Resting state data

The localization results for resting state data analysis from six subjects are shown in Figure 7. For resting state analysis, there is no clear baseline data for background noise covariance estimation. As is seen, Champagne with CB_NL can localize all subjects’ brain activity near the midline occipital lobe or posterior cingulate gyrus during rest. For resting state analysis, even though there is no pre-stimulus data for background noise estimation, Champagne with CB_NL is able to learn the underlying noise and still recovers reasonable activity.

**Figure 7:**
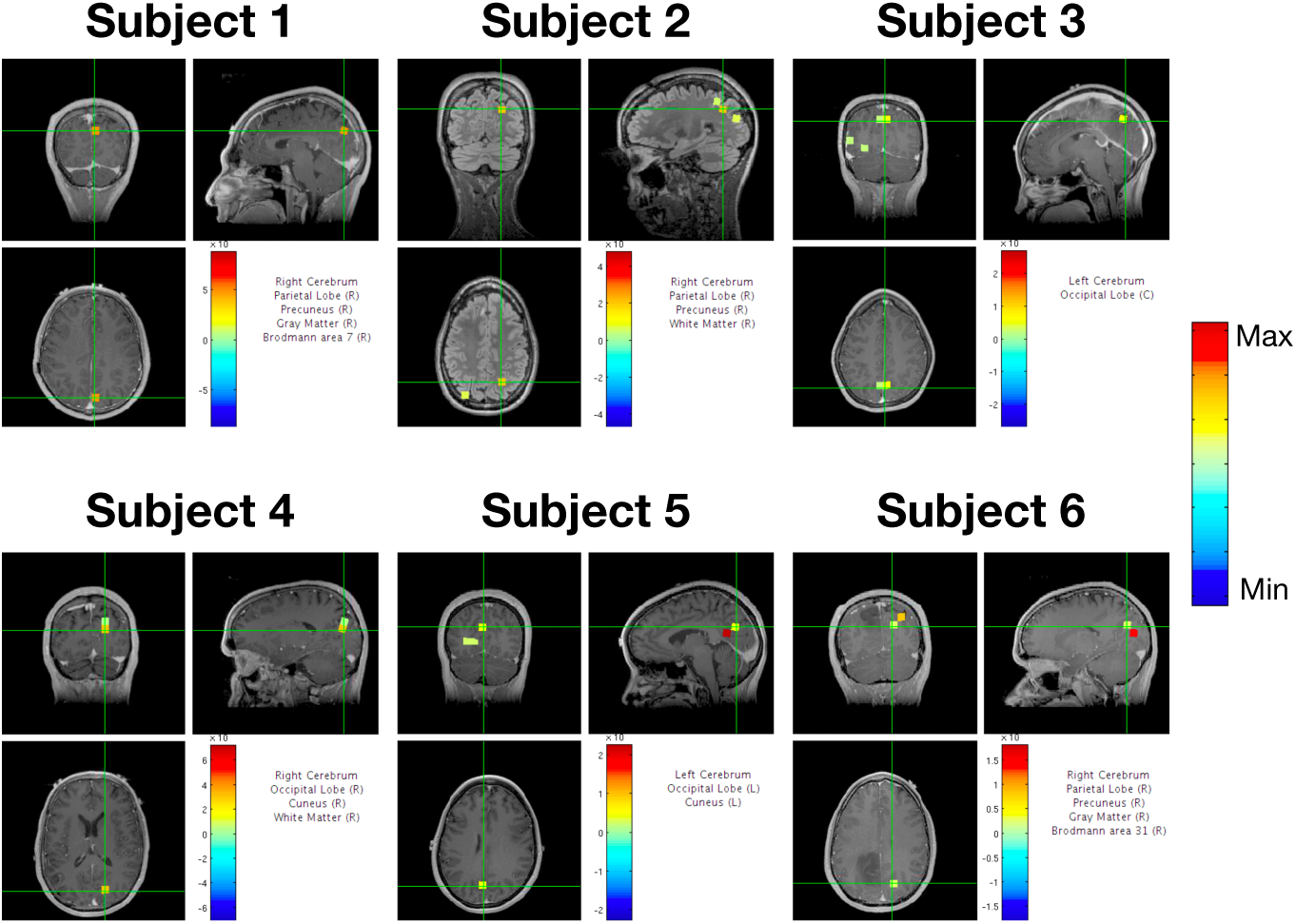
Localization results of spontaneous brain activity for six subjects using Champagne with CB_NL.

In summary, from the reconstructed results of our real data sets, Champagne with noise learning consistently produces better results than Champagne with Noise_Sub (0.1% to 10%) and sLORETA. Champagne with CB_NL is able to localize brain activity accurately with a few trials or even a single trial, while Champagne with Noise_Sub (0.1% to 10%) fails to localize when the number of trials is less than 12. Even though pre-stimulus data is unavailable for resting state analysis, Champagne with CB_NL is able to learn the underlying noise and still recover reasonable activity.

## 5. Discussion

This paper derives several robust ways to estimate contributions to sensors from noise without the need for additional “baseline” or “control” data, while preserving robust reconstruction of complex brain source activity and performance features of the sparse source reconstruction algorithm Champagne. The underlying data estimation portion of the algorithms are based on a principled cost function which maximizes the marginal likelihood or a convex lower bound on the marginal likelihood of the data, resulting in fast and convergent update rules.

In our novel algorithms, we further optimize the cost function using several robust algorithms to learn the noise. Noise learning is accomplished by making assumptions about the structure of the noise followed by Bayesian inference to derive update rules for noise estimation. Scalar covariance matrix can be used for the modeling of homoscedastic noise, while diagonal covariance matrix can be used to model heteroscedastic noise. In our novel noise estimation algorithms, we employ the following strategies for updating the noise covariance: Marginal Likelihood (ML) maximization, expectation maximization (EM), and convex bounding (CB) based Bayesian inference. The new algorithms readily handle a variety of configurations of point brain sources under high noise and interference conditions without the need for additional “baseline” or “control” measurements - a situation that commonly arises in resting state data analysis. Computationally, it can be shown that augmenting the voxel leadfield matrix with an identity matrix corresponding to sensor noise can be used to simultaneously update the voxel and noise covariances using convex-bounding methods efficiently.

The novel algorithms display significant theoretical and empirical advantages over the existing benchmark Champagne algorithm when the noise covariance cannot be accurately determined in advance. Simulations were developed to explore noise learning algorithmic performance for complex source configurations with highly correlated time-courses, multiple point sources, and high levels of noise and interference. These simulations demonstrated that noise-learning Champagne algorithms outperform traditional Champagne with an incorrect noise covariance as they show higher *AP* scores. Furthermore, noise-learning Champagne’s performance demonstrates that noise learning is robust even when the algorithms are initialized to incorrect noise values.

In general, it is difficult to evaluate localization algorithm performance with real data since the ground truth is unknown. For this reason, we chose real data sets that have well-established patterns of brain activity (SEF and AEF). We also demonstrate that even though pre-stimulus data for resting state analysis does not exist, our novel algorithm is able to learn the underlying noise and still recover reasonable activity. Performance on these real data sets demonstrates that Champagne with noise learning is superior in localizing real brain activity when compared to Champagne with Noise_Sub and sLORETA.

Since brain activity has a very low signal-to-noise ratio compared to background activity, many trials are often required for reconstruction of evoked fields. Using our novel noise-learning algorithm, we are able to robustly localize brain activity with a few trials or even with a single trial in our SEF and AEF datasets, which is a revolutionary improvement in electromagnetic brain imaging. In fact, data collection times may be dramatically reduced up to ten-fold, which is particularly important in studies involving children with autism, patients with dementia, or any other subjects who have difficulty tolerating long periods of data collection.

We now discuss related work on noise covariance estimation for electromagnetic imaging. Cross-validation and likelihood estimation methods have been proposed by Engemann *et al*. with the availability of separate baseline data for noise covariance estimation [21, 22]. More complex spatiotemporal noise covariance structures have also been estimated from separate baseline measurements [23, 24, 25, 26]. Joint estimation of brain source activity and noise covariance have been previously proposed for Type-1 penalized likelihood methods. Massias *et al*. [27] proposed a Smoothed Generalized Concomitant Lasso (SGCL) algorithm, which examined mixed-norm optimization for MEG imaging. This model includes a single scalar parameter for noise, independent of the multiple noise levels present in heterogeneous data. Subsequently, Quentin *et al*. extended the SGCL framework to a Concomitant Lasso with Repetitions (CLaR) estimator [28] that can cope with more complex noise structure estimated from non-averaged measurements. In contrast to these Type-I likelihood estimation methods, the present Champagne algorithm with noise learning uses Type-II likelihood estimation methods. We have previously shown that Type-I likelihood methods which make use of non-factorial, lead-field and noise-dependent priors can be shown to have a dual with Type-II likelihood methods with comparable performance [29]. However, in general, in our previous papers [20], performance of Type-II likelihood estimation methods yields superior results to Type-I methods with factorial priors.

Here, we assume noise is independent between different sensors in order to make estimating source variance and noise covariance simultaneously a tractable problem. In the future, if baseline data is available and can be used to estimate the gain matrix of interference and noise, adding other noise structures into the forward model, such as low-rank Toeplitz noise, is hoped to improve the learning of dependent or correlated sensor noise and to further improve the estimation of brain electromagnetic activity in noisy environments. In addition, the application of methods in this paper can also be derived by augmenting the leadfield matrix with an identity matrix, see Appendix D, which will be further detailed and tested as part of our future work. Extending the current Gaussian noise priors to more realistic non-Gaussian priors may also significantly improve the results, which will also be part of our work in the future. Using a noise covariance model based on a single Kronecker product of spatial and temporal covariance in the spatiotemporal analysis of MEG data has been demonstrated to provide improvement in the results over that of the commonly used diagonal noise covariance model [23, 24, 25, 26], which will be a potential extension for Champagne algorithm to improve the accuracy of brain source activity and noise estimation.

## Acknowledgment

The authors would like to thank Danielle Mizuiri and Anne Findlay for collecting much of the MEG data, and all members and collaborators of the Biomagnetic Imaging Laboratory for their support. This work was supported by NIH grants R01EB022717, R01DC013979, R01NS100440, R01DC176960, R01DC017091, R01AG062196 UCOP-MRP-17-454755, and T32EB001631 from the NIBIB, and by the European Research Council (ERC) under the European Union’s Horizon 2020 research and innovation programme (Grant agreement No. 758985). AH acknowledges scholarship support from the Machine Learning/Intelligent Data Analysis research group at Technische Universität Berlin and partial support from the Berlin International Graduate School in Model and Simulation based Research (BIMoS), the Berlin Mathematical School (BMS), and the Berlin Mathematics Research Center MATH+.

## Appendix A. Learning noise covariance using marginal likelihood maximization

The first and direct way to estimate the noise covariance **Λ** is setting the derivative of the cost function with respect to **Λ** to zero and fixing ***α***,

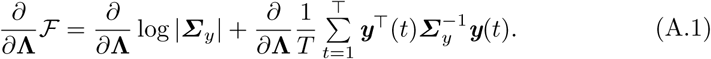

Using ***Σ***_*y*_ = **Λ** + ***LαL***^*T*^ with the following matrix lemma,

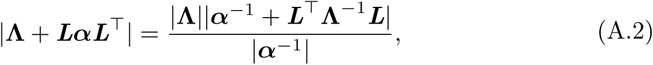

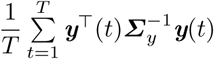 from the second term in the right-hand side of Eq. (A.1) can be computed as

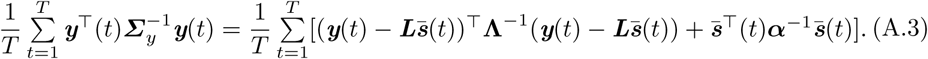

Since **Λ** = diag(λ_1_, λ_2_, …, λ_*M*_), the update rule for λ_*M*_ is expressed as:

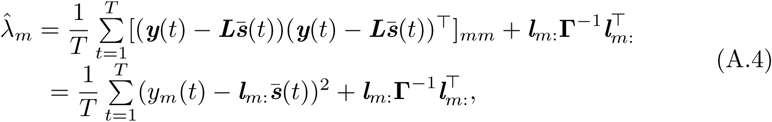

where **Γ** = ***α***^−1^ + ***L*^*T*^Λ^−1^*L***. An alternative derivation of noise covariance using the Expectation Maximization algorithm is shown below, which results in identical update rules.

## Appendix B. Leaning noise covariance using Expectation Maximization (EM) algorithm

The second way to estimate the noise covariance **Λ** is using EM algorithm with the following cost function [2] expressed as

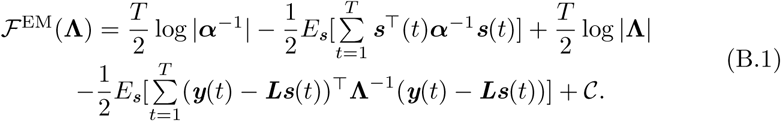

where 𝒞 expresses terms that do not contain **Λ**. Setting the derivative of the cost function Eq. (B.1) with respect to *λ*_*m*_ to zero generates the update rule for noise covariance as follows

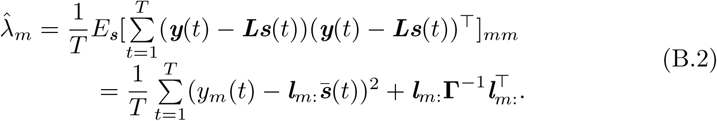

The EM update rule for noise covariance in the above equation is the same as the update rule derived by direct maximizing marginal likelihood.

## Appendix C. Learning noise covariance using convex bounding approach

The third way to estimate the noise covariance is using an auxiliary cost function [2] which is based on the convex bounding approach [10]. Noise covariance update rules can be derived through convex bounding of the marginal likelihood,

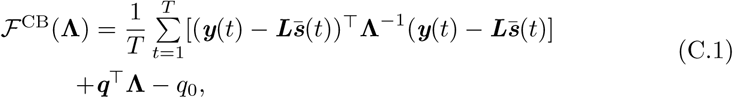

where ***q*** = [*q*_1_, *q*_2_, …, *q*_*M*_]^*T*^ is an auxiliary variable, and *q*_0_ is a scalar term. For the *n*-th sensor, the convex bounding update rule for noise variance of the *n*-th sensor is obtained by setting the derivatives of ℱ^CB^(**Λ**) with respect to *λ*_*m*_ to zero, resulting in

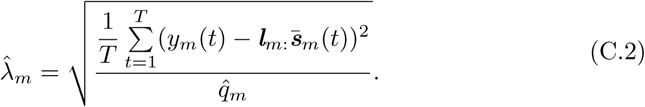

The update rule for ***q*** is equivalent to finding a hyperplane ***q*^*T*^Λ** − *q*_0_ that forms a closest upper bound of log |***Σ***_*y*_|. Such a hyperplane is found as the plane that is tangential to log |***Σ***_*y*_| [1]. Therefore, the updated value 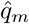 is given by

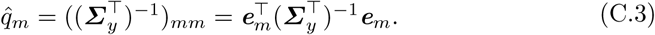

where ***e***_*m*_ is a *M* × 1 vector which is the leadfiled matrix for *m*-th sensor’s noise, where the element in *m*-th row is 1, the others are 0.

## Appendix D. Estimating brain sources and noise covariance simultaneously using convex-bounding based approach

We can estimate brain source activity and noise variance simultaneously with slight modifications to the generative model. We assume that each sensor measurement is the summation of whole brain activity and one noise source. The leadfield matrix for the noise activity is assumed as an *M* × *M* identity matrix ***I*** = [***e***_1_, …, ***e***_*M*_]. The generative model can then be rewritten as,

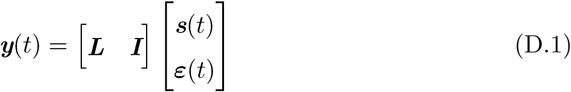

where

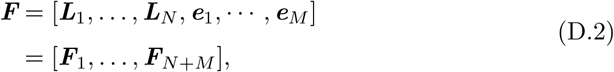

and

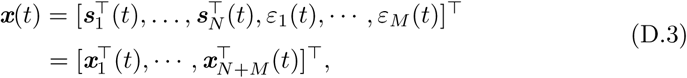

are the augmented leadfield matrix ***F***_*k*_ and the time courses for the brain activity and noise ***x***_*k*_(*t*), (*k* = 1, …, *N* for sources, *k* = *N* +1, …, *N* +*M* for noise). We also define the augmented prior hyperparameters ***ν*** for source and noise activity as

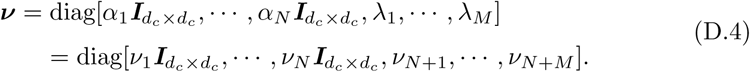

Utilizing the convex bounding on the marginal likelihood results in fast convergence properties in the following update rules:

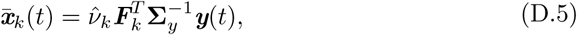

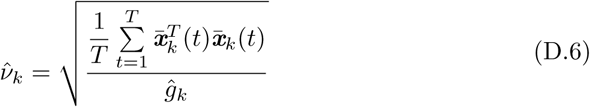

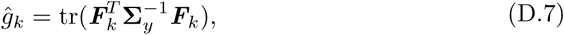

where, *g*_*k*_ is an auxiliary variable. ***Σ***_*y*_ = ***FνF***^*T*^ is the model data covariance matrix. In summary, the augmentation algorithm simultaneously estimates brain sources and noise activity 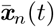 by iterating between Eqs. (D.5), (D.6) and (D.7).

